# Spatial Multiomics Reveal the Role of Wnt Modulator, Dkk2, in Palatogenesis

**DOI:** 10.1101/2023.05.16.541037

**Authors:** Jeremie Oliver Piña, Resmi Raju, Daniela M. Roth, Emma Wentworth Winchester, Cameron Padilla, James Iben, Fabio R. Faucz, Justin L. Cotney, Rena N. D’Souza

**Affiliations:** Section on Craniofacial Genetic Disorders, Eunice Kennedy Shriver National Institute of Child Health and Human Development, National Institutes of Health, Bethesda, MD, USA; Department of Biomedical Engineering, University of Utah, Salt Lake City, UT, USA; School of Dentistry, University of Alberta, Edmonton, AB, CA; University of Connecticut School of Dental Medicine, Farmington, CT, USA; Molecular Genomics Core, Eunice Kennedy Shriver National Institute of Child Health and Human Development, National Institutes of Health, Bethesda, MD, USA; Department of Genetics and Genome Sciences, University of Connecticut School of Medicine, Farmington, CT, USA

**Keywords:** Cleft Palate, Signal Transduction, Craniofacial Anomalies, Craniofacial Biology/Genetics, Gene Expression, Single Cell Sequencing

## Abstract

Multiple genetic and environmental etiologies contribute to the pathogenesis of cleft palate, which constitutes the most common among the inherited disorders of the craniofacial complex. Insights into the molecular mechanisms regulating osteogenic differentiation and patterning in the palate during embryogenesis are limited and needed for the development of innovative diagnostics and cures. This study utilized the *Pax9^-/-^* mouse model with a consistent phenotype of cleft secondary palate to investigate the role of *Pax9* in the process of palatal osteogenesis. While prior research had identified upregulation of Wnt pathway modulators *Dkk1* and *Dkk2* in *Pax9^-/-^* palate mesenchyme, limitations of spatial resolution and technology restricted a more robust analysis. Here, data from single-nucleus transcriptomics and chromatin accessibility assays validated by *in situ* highly multiplex targeted single-cell spatial profiling technology suggest a distinct relationship between *Pax9+* and osteogenic populations. Loss of *Pax9* results in spatially restricted osteogenic domains bounded by *Dkk2*, which normally interfaces with *Pax9* in the mesenchyme. These results suggest that Pax9-dependent Wnt signaling modulators influence osteogenic programming during palate formation, potentially contributing to the observed cleft palate phenotype.

## Introduction

The Wnt/β-catenin signaling pathway has an important regulatory role in the development and homeostasis of the skeleton (Huybrechts et al. 2020; Joeng et al. 2017; Krishnan et al. 2006), including the craniofacial complex. We and others have shown that Wnt signaling pathway genes play crucial roles in palatogenesis, specifically driving mesenchymal cell proliferation and differentiation (Jia et al. 2017a; Jia et al. 2017b; Li et al. 2017; Jia et al. 2020; Janeckova et al. 2023; Pina et al. 2023). Intracellular mediators of canonical Wnt signaling, such as β-catenin, promote lineage specification of mesenchymal stem cells and the differentiation of osteogenic precursors into functional osteoblasts and osteocytes (Glass et al. 2005; Cruciat and Niehrs 2013; Kim et al. 2013). Notably, mutations in coding sequences of Wnt factors result in skeletal dysplasias with clinical phenotypes that involve isolated alterations in bone mass (Regard et al. 2011; Martinez-Gil et al. 2022). Genome-wide association studies (GWAS) have implicated *PAX9*, the paired homeodomain-containing transcription factor, in craniofacial morphogenesis (Shaffer et al. 2016), hinting at the plausibility of genetic variants in this region influencing normal facial variation.

*Pax9^-/-^* mice offer a valuable resource for studies on palate development as they consistently demonstrate defects in elongation, delayed elevation, failed contact, and fusion of the palatal shelves (Peters et al. 1998; Jia et al. 2017a). We have used *Pax9^-/-^*mice to demonstrate a unique molecular relationship between the Wnt signaling pathway and Pax9 within the posterior domain of the developing palate (Jia et al. 2017a; Jia et al. 2020). Our results suggest that Pax9 shares a functional relationship with Dkk1 and Dkk2, specific modulators of Wnt signaling during secondary palate development (Jia et al. 2020). It is known that the Dkk1 and Dkk2 proteins block the binding of Wnt ligands to the low-density lipoprotein receptor-related protein (LRP) 5 and 6 at the cell surface (Semenov et al. 2001). This competitive inhibition of Wnt effector molecule function directly antagonizes the activation of Wnt signal transduction (Tamai et al. 2000). Further evidence of the Dkk’s role in Wnt signaling modulation is evidence in our prior work, wherein the genetic reduction of *Dkk1* during palatogenesis corrected secondary palatal clefts in *Pax9*^-/-^ mice with restoration of Wnt signaling activities (Jia et al. 2017; Jia et al. 2020). Furthermore, ChIP-qPCR assays showed that Pax9 directly binds to regions near the transcription start sites of *Dkk1* and *Dkk2* (Jia et al. 2020). Thus, we concluded that the molecular mechanisms underlying Pax9’s role in modulating Wnt signaling activity during palate development is likely to involve the inhibition of *Dkk1* and *Dkk2* expression.

The advancement of single-cell and spatial transcriptomics technologies provides us with valuable tools to further explore the relationship between Pax9 and Wnt signaling. Here, we demonstrate that the global loss of *Pax9* disrupted the spatial expression domains of key Wnt modulator genes – in particular, *Dkk2* – at critical phases of secondary palate formation. We present a Wnt-centered framework in high spatial resolution for osteogenic patterning, the balance of which depends on the regulatory influence of *Pax9*. This study expands contemporary understanding of the functional roles played by *Pax9* and its overlapping Wnt modulators to bring about differentiation and patterning morphogenesis of the secondary palate.

## Materials and Methods

### Animals

All animal procedures were approved by the National Institutes of Health, National Institute of Child Health and Human Development Animal Care and Use Committee (ACUC), under Animal Study Protocol (ASP) #21-031. C57BL/6J mice were obtained from the Jackson Laboratory. Inbred strain of female C57BL/6 mice were utilized for all experiments. *Pax9*^+/−^ mice were provided by Dr. Rulang Jiang (Cincinnati Children’s Hospital) and generated as described previously (Zhou et al. 2013). Timed pregnancies were conducted via vaginal plug identification, with day 0.5 indicating date of identification. Genotyping of all mice included in this study was performed by Transnetyx, Inc. This study conforms to the ARRIVE 2.0 guidelines.

### Single-nucleus RNA + ATAC Sequencing (Multiome-RNA+ATAC-seq)

In this study, we generated a new single-nucleus gene (snRNA) and transposase-accessible chromatin (snATAC) Multiome dataset of microdissected *Pax9*^-/-^ secondary palate tissue from embryonic day (E) 13.5 (n=3 pooled biological replicates) for direct comparison to our previously generated Multiome dataset on normal secondary palate tissue from E13.5 and E15.5 mouse embryos (Pina et al. 2023). To properly compare the new Multiome dataset to those generated in our prior study, all methodological steps were replicated as described previously (Pina et al. 2023).

### Single-nucleus RNA + ATAC-seq Bioinformatics Analysis

Raw fastqs were aligned to mm10 genome build using the standard Cellranger multiomics settings and imported to ArchR (v.1.0.1). Doublets were identified and filtered, and cells were filtered for minimum 4 TSS enrichment, 2500 fragments per cell. Dimensionality reduction was performed using LSI based on the cell by fragment matrix and cell by gene matrix, and clusters were identified. Peaks were called based on original clustering, discarding reads from promoters (2500bp +/-TSS) and exons. Cluster assignments were confirmed using canonical marker genes based on gene expression, and gene ontology enrichment of marker genes identified using getMarkerFeatures function of ArchR. Statistical significance and strength of enrichments were determined using t-test, grouping cells by cluster. Detailed scripts and bioinformatic methodology of snRNA-seq analysis can be found on the Cotney Lab GitHub (https://github.com/emmawwinchester/mousepalate).

### Micro-Computed Tomography (*µ*CT) 3D Reconstruction and Segmentation

Embryonic day 15.5 heads (n=3 biological replicates per genotype) fixed overnight in 10% NBF were scanned using a ScanCo µCT 50 *ex vivo* cabinet system (Scanco Medical, Brüttisellen, Switzerland) in 70% ethanol using the following parameters: 70kVp, 85µA, 0.5mm AI filter, 900-ms integration time, and 10µm voxel size. 3D rendering and segmentation of reconstructed files were performed using 3D Slicer software (slicer.org) using the semi-automatic segmentation function ‘Islands’, with standardized thresholding and island size.

### Xenium In Situ mRNA Localization and Analysis

Xenium *in situ* sub-cellular mRNA detection technology was employed as previously reported following all manufacturer guidelines and specifications (Janesick et al. 2022). We custom designed a 350-plex targeted gene panel (Supplementary File) to detect mRNA expression for cell type identification as well as signaling pathway interactions selected and curated primarily based on single cell atlas data we generated in our previous work on palate development (Pina et al. 2023). In brief, Xenium *in situ* technology allows for probes to be designed to contain two complementary sequences that hybridize to the target RNA as well as a third region encoding a gene-specific barcode, so that the paired ends of the probe bind to the target RNA and ligate to generate a circular DNA probe. If the probe experiences an off-target binding event, ligation should not occur, suppressing off-target signals and ensuring high specificity. For this assay, we formalin fixed, and paraffin embedded 3 biological replicates of *Pax9*^-/-^ and 3 biological replicates of *Pax9*^+/+^ mouse embryos at embryonic day 14.5. Whole heads were processed, embedded, and sectioned to the level of the 1^st^ molar. All six samples were assayed in a single batch under identical conditions. The post-Xenium H&E staining followed Demonstrated Protocol CG000160 from 10x Genomics, Inc. All analyses were generated and exported from the 10x Genomics open-access software, Xenium Explorer v1.2, following previously published guidelines from the manufacturer. Regions of interest were defined and standardized between sections analyzed, as were number of cells per frame. The differential expression analysis seeks to find, for each cluster, features that are more highly expressed in that cluster relative to the rest of the sample. A differential expression test was performed between each cluster and the rest of the sample for each feature. The Log_2_ fold-change (L2FC) is an estimate of the log_2_ ratio of expression in a cluster to that in all other cells. A value of 1.0 indicates 2-fold greater expression in the cluster of interest. The p-value is a measure of the statistical significance of the expression difference and is based on a negative binomial test. The p-value reported here has been adjusted for multiple testing via the Benjamini-Hochberg procedure. Features were filtered (Mean object counts > 1.0) and retained the top N features by L2FC for each cluster.

## Results

### A Pax9-Enriched Sub-Cluster is Transcriptionally Distinct from Other Mesenchymal Cell Populations in the Developing Palate

First, we further analyzed our recently published datasets assessing cell type-specific gene expression and binding motif enrichment using integrated snRNA- and snATAC-sequencing of micro dissected *Pax9*^+/+^ secondary palate tissue at E13.5 and E15.5 of murine development (Pina et al. 2023). Initial clustering and gene ontology enrichment analysis of marker genes from these populations determined the presence of six general cell types: epithelium, mesenchyme, muscle cells, neural cells, endothelium, and blood cells (Fig. 1A). Further isolated analysis of the mesenchyme identified eight subtypes of this population. Canonical marker genes separate these cells into four broad categories: osteogenic cells, chondrogenic cells, generalized mesenchyme, and *Pax9+* mesenchyme (Pina et al. 2023).

**Figure 1.**
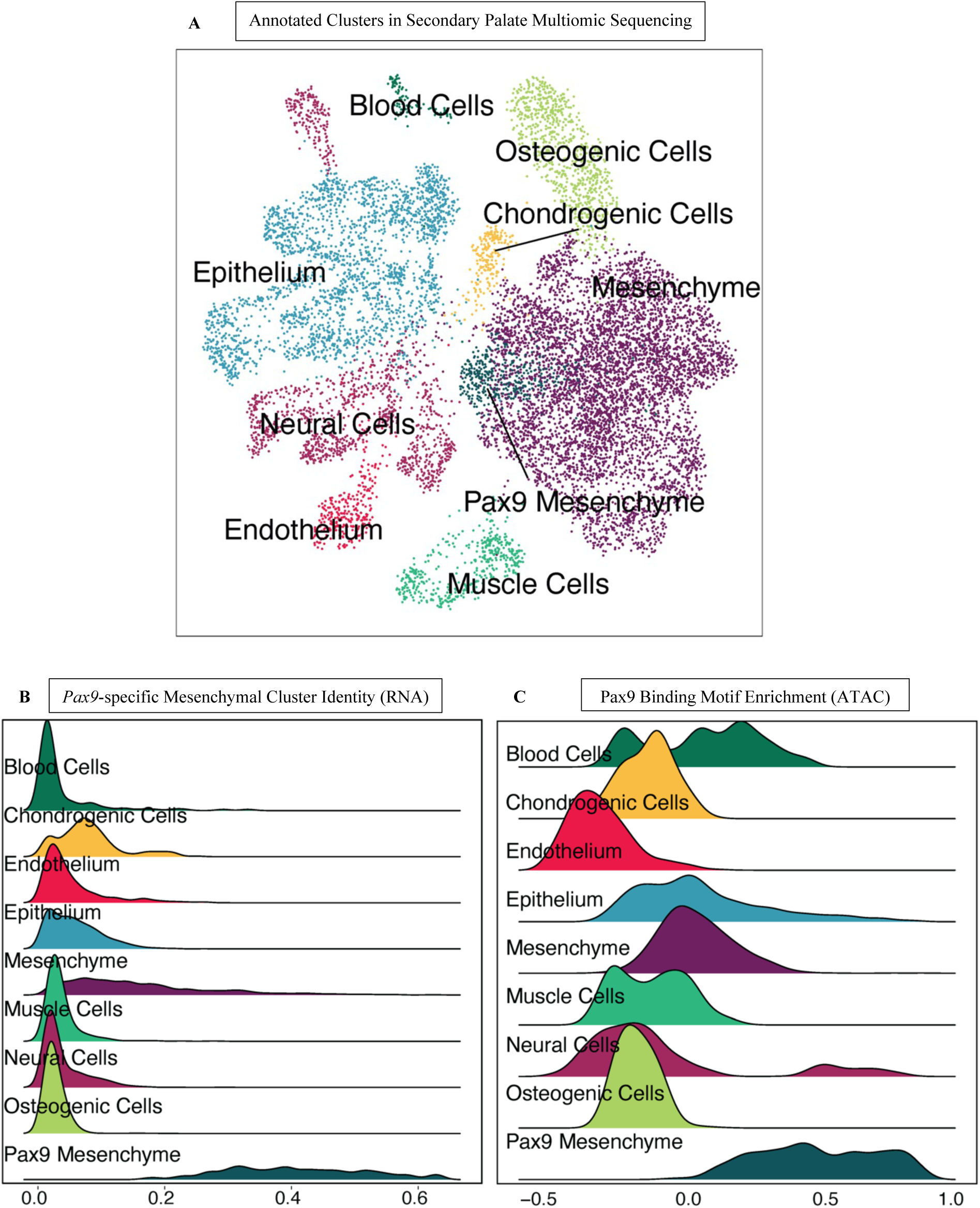
Single-nucleus Multiomic sequencing (snRNA+ATAC-seq) of normal secondary palate identifies *Pax9*-enriched cell cluster distinct from other mesenchymal populations. A Uniform manifold approximation projection (UMAP) of Initial clustering and gene ontology enrichment analysis of marker genes from pooled biological triplicate microdissected secondary palate tissues at embryonic days 13.5 and 15.5. **B** Enrichment of Pax9 expression in *Pax9*+ Mesenchyme compared to all other cell types in the secondary palate snRNA-seq dataset. **C** Bias of Pax9 binding motif in regions of chromatin uniquely accessible in *Pax9*+ Mesenchyme from snATAC-seq dataset.

When compared to all cell types in the developing palate, we observed that the population of *Pax9+* mesenchymal cells demonstrated a 3.62 log_2_-fold enrichment of *Pax9* expression (FDR 3.01e-04) (Fig. 1B). Similarly, we noted a bias of the *Pax9* motif (Castro-Mondragon et al., 2023) within peaks accessible within the *Pax9* mesenchyme cluster, indicating bias of the *Pax9* motif within regions of accessible chromatin open in this population (Fig. 1C). Notably, chromatin regions accessible in the osteogenic cell population were depleted for the *Pax9* motif and demonstrated minimal *Pax9* expression. Hence, *Pax9* may influence osteogenic programming via Wnt signaling modulation of adjacent cell types within the developing palate mesenchyme which exert paracrine effects on neighboring osteogenic cells.

### Loss of Pax9 Alters Wnt Modulator Expression and the Growth of Osteogenic Zones in the Midline of Palatal Shelves Extension

Next, we sought to understand the effects of global loss of *Pax9* on the development of the secondary palate osteogenic-enriched population. µCT revealed three-dimensional insufficiency of the mesial ossification front of the palatine bone at E15.5, despite indications of mature bone (Fig. 2A). Coronal cross-sectional analysis exhibited dense, spongy bone in *Pax9*^-/-^ embryos (Fig. 2A). To better assess the downstream molecular consequences of the global loss of Pax9 expression in the secondary palate, we generated a new Multiome-seq dataset of murine E13.5 *Pax9*^-/-^ palatal shelves to directly compare to our previously generated dataset of normal secondary palate development at the same time point. Integrated Multiome-seq analysis of *Pax9*^+/+^ and *Pax9*^-/-^ datasets revealed a relatively greater proportion of early and late osteoprogenitor cells in E13.5 *Pax9*^-/-^ secondary palate compared to the E13.5 *Pax9*^+/+^ secondary palate samples (Fig. 2B). Furthermore, the expression domain of key Wnt signaling modulators, *Dkk1* and *Dkk2*, was notably expanded in the *Pax9^-/-^* secondary palate (Fig. 2C). Together, these morphological and molecular features of the *Pax9*^-/-^ secondary palate could indicate a disruption in Wnt signaling dynamics – evidenced by the relative increase in expression of direct Wnt antagonists – which correlates to an alteration of palatal ossification processes. This may suggest a critical role for Pax9-dependent Wnt signaling in maintaining proper palatal bone formation during development.

**Figure 2.**
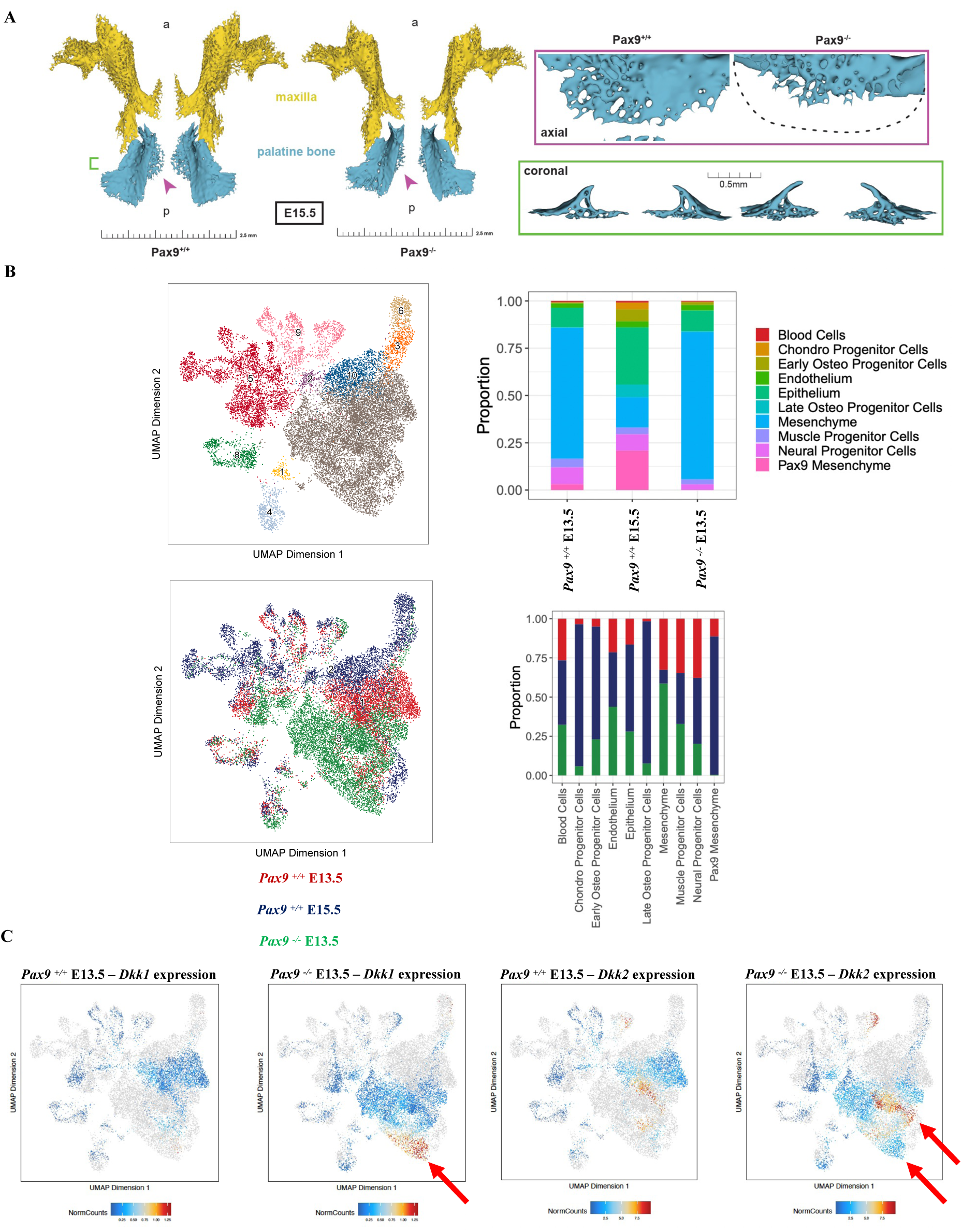
Loss of global Pax9 results in failure of palatal bone outgrowth and midline fusion, with notable increase in Wnt modulator and osteoprogenitor cell expression. **A** Micro-computed tomography 3D reconstruction of paired maxillary (yellow) and palatine bones (blue) from *Pax9*+/+ and *Pax9*^-/-^ biological triplicates at E15.5, presented from a superior axial view (top) and coronal orthogonal slice (bottom). Dotted outline represents normal palatine bone midline extension silhouette superimposed over *Pax9*-/-palatine bone; purple arrowheads highlight osteogenic fronts; green bracket outlines regional of palatine bone coronal 3D view. **B** UMAP projections and relative proportions of cell types between groups in *Pax9*^+/+^ and *Pax9*^-/-^ secondary palate Multiome datasets highlights heterogeneous transcriptomic signatures, with a slight relative increase in early and late osteoprogenitor cell markers in the *Pax9*-/-compared to the *Pax9*+/+ at E13.5 (n=3 pooled, microdissected secondary palate tissue from *Pax9*-/-embryos at E13.5 in biological triplicate, as with *Pax9*^+/+^ dataset for equal comparison). **C** UMAP plots of Wnt modulators *Dkk1* and *Dkk2* at E13.5 in both *Pax9*^+/+^ and *Pax9*^-/-^ secondary palate reveal increased normalized counts specifically within mesenchymal clusters in the absence of Pax9.

### Spatial Dysregulation of Wnt Effector and Modulator Genes in Pax9^-/-^ Cleft Secondary Palate

To study whether *Pax9* regulates the commitment of palatal mesenchymal progenitor cells to the osteogenic lineage through Wnt signaling, we performed highly multiplex targeted single cell spatial profiling analyses (Xenium *in situ*, 10x Genomics, Inc.). Age-matched (E14.5) whole embryonic heads from *Pax9*^+/+^ and *Pax9*^-/-^ murine litters (n=3 biological replicates per group) were included. As the first application of such spatial transcriptomic technology in developmental biology, we custom designed a panel of 350 unique marker gene probes (Supplemental Table 1) to dissect differential cell types and effector-ligand signaling interactions within the secondary palate with and without the functional presence of Pax9.

Spatial transcriptomic analysis revealed 28 different cell clusters in both *Pax9*^+/+^ and *Pax9*^-/-^ samples (n=3 biological replicates per genotype were run and analyzed) (Fig. 3A-B). Transcriptomic signature differences were identified in differential expression analysis between *Pax9*^+/+^ and *Pax9*^-/-^ samples. Clusters in the secondary palate region of interest were identified based on spatial localization within the coronal sections, and genes were analyzed within these clusters specifically. Clusters with enriched genes related to epithelium, osteogenic mesenchyme, extracellular matrix, and ciliated cells (both epithelium and mesenchyme) were identified for each sample analyzed in each group (Fig. 3C). Notably, there were two distinct clusters (1, 7) identified in the *Pax9*^+/+^ palate samples related to epithelial cell marker gene expression, while there was only one distinct cluster (7) present in the *Pax9*^-/-^ samples. We noted that epithelial expression of *Wnt5a* and *Wnt7b* was present in abundance in the *Pax9*^+/+^ palate epithelium, while neither was found to be enriched in cluster 7 of the *Pax9*^-/-^ samples. In the osteogenic mesenchyme-associated clusters, the *Pax9*^-/-^ palate demonstrated five (2, 8, 9, 10, 12) unique clusters of enriched genes, compared to just four (2, 3, 6, 10) clusters in the *Pax9*^+/+^ palate. Notably, Wnt signaling modulators, such as *Sfrp2* differentially expressed in clusters 2 and 9, were identified to be enriched in the *Pax9^-/-^* palate osteogenic mesenchyme in the *Pax9*^+/+^ palate samples. Similarly, the extracellular matrix clusters revealed enhanced Wnt modulator expression as well as the appearance of expanded defining marker genes in the absence of Pax9, including the early osteoprogenitor cell marker, *Crabp1*. Finally, in the primary cilia-associated clusters, several genes showed a differential down regulation in the absence of Pax9, including *Dynlrb2, Foxj1, Capsl, Mapk13, Krt7, Tuba1b*, and *Wnt7b*. Taken together, across all cell types identified within the secondary palate, our in-situ analysis provides additional evidence of a concordant genetic relationship between Pax9 and Wnt signaling during palate development (Fig. 3D).

**Figure 3.**
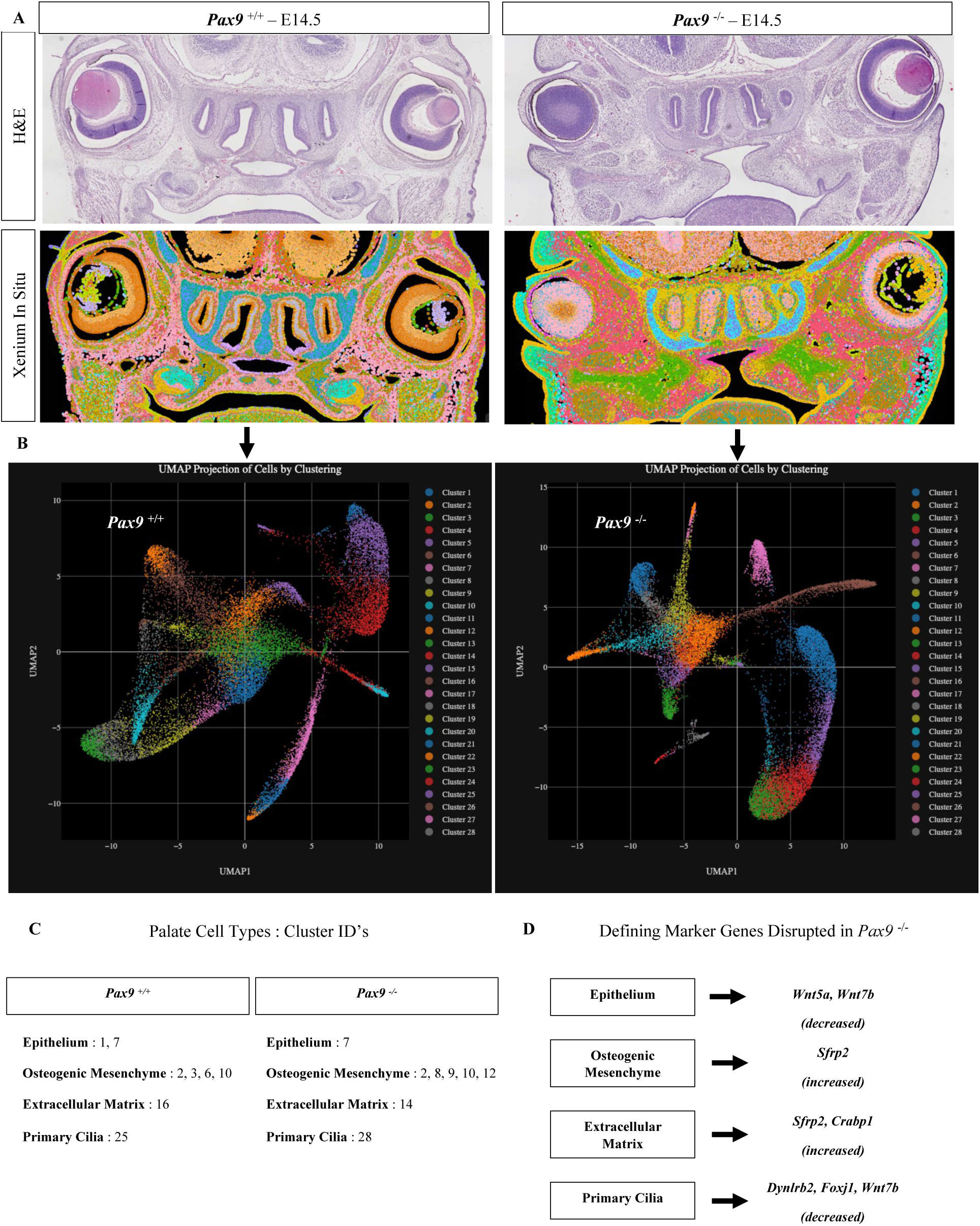
Highly multiplex in situ mRNA localization in normal and *Pax9-/-* palatal shelves reveal disruption of spatial patterning and mesenchymal progenitor cell differentiation. Whole embryonic heads at E14.5 of *Pax9*^+/+^ and *Pax9*^-/-^ (n=3 biological replicates of each) were formalin-fixed and coronally paraffin-embedded for H&E and Xenium in situ spatial transcriptomic analysis. **B** UMAP plots of all 28 clusters identified in Xenium in situ assay for each sample. **C** Defining palate cell types and respective cluster ID’s based on Xenium custom panel marker genes explored (see Supplemental File). **D** Specific cluster and gene changes between *Pax9*^+/+^ and *Pax9*^-/-^ samples reveal Wnt signaling dysregulation across cell types.

### Palatal Bone in Pax9^-/-^ Cleft is Bounded by Dkk2

Validating and spatially resolving our observations from the Multiome-seq analysis, we observed a notable increase in the number of *Dkk1* and *Dkk2* transcripts localized within the *Pax9*^-/-^ cleft palate mesenchyme (Fig. 4A). Interestingly, *Dkk1*’s expression domain is localized primarily within the osteogenic zones, while *Dkk2*’s expression bounds the mesial border of midline osteogenic extension of the palatine bone (Fig. 4B). Furthermore, we noted spatial alteration in expression domains of *Fgfr1*, *Gli1*, *Bmp2*, and *Bmp4* in the *Pax9^-/-^* palatal shelves, notably in the mesial aspect of the elevated shelves, the domain of *Pax9* enriched expression in normal palatal shelves (Supplemental Fig. 1, Tables 2 and 3). This higher spatial resolution of specific mesenchymal compartment gene enrichment extends and enlightens our prior studies on these molecules *in situ*. Specifically, the mesial enrichment of *Dkk2* (alongside overlapping pathway effector deficiencies) may hint at its functional role within the *Pax9^-/-^* cleft palate, actively antagonizing Wnt-driven osteogenic extension to the midline.

**Figure 4.**
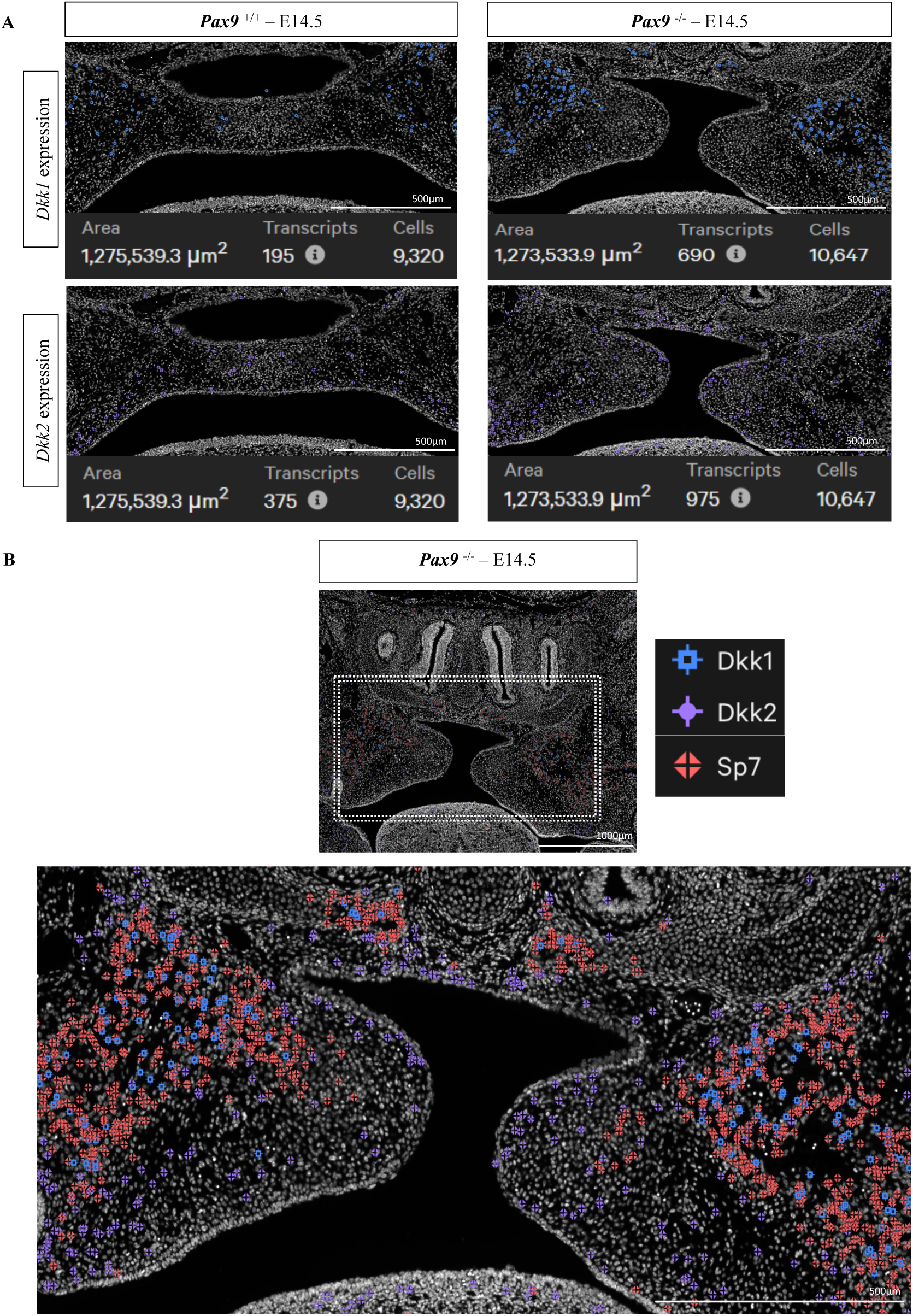
Spatial dysregulation of Wnt modulator expression within *Pax9-/-* palatal shelves. **A** Quantitative and spatial disruption of both *Dkk1* and *Dkk2*, with notable increase of *Dkk2* in the absence of Pax9; Region of interest was selected using the Xenium Analyzer v1.2 with square area shown. Transcripts detected for each gene were compared within the same square ROI and quantified automatically within the software program. All quantitation performed in biological triplicate, with representative sections shown in this image panel. **B** *Dkk2* enrichment bounds the mesial border of midline osteogenic extension of the palatine bone, marked by *Sp7* transcripts, in *Pax9*-/-cleft palate.

## Discussion

Pax9 is a master orchestrator of patterning and morphogenesis throughout the body – from the thymus, ultimobranchial bodies, limb bud, and tooth organ, to the axial skeleton (Peters et al. 1998; Peters et al. 1999). This highly conserved transcription factor has been shown to functionally interact with a number of mesenchymal factors and signaling pathways required for palate development to ensue, which act in concert to regulate cell proliferation and differentiation of cranial neural crest-derived mesectoderm (Neubuser et al. 1997; Hilliard et al. 2005; Ichikawa et al. 2006; Jia et al. 2017a; Li et al. 2017; Jia et al. 2020). The role of Pax9 in patterning during development of multiple organ systems, including the palate, are clearly established. However, prior to the present study, this has not been explored sufficiently in the context of palate osteogenesis.

The resultant cleft palate phenotypes in the setting of either premature (Mori-Akiyama et al. 2003; Sweat et al. 2020) or delayed (Baek et al. 2011) palate osteogenesis exemplify the importance of precise temporal onset and spatial regulation of intramembranous ossification. Intriguingly, and supportive of our hypothesis that Wnt-mediated osteogenic programming is disrupted in the absence of Pax9, a premature ossification and cleft palate phenotype is described in *Sox9*^-/-^ (Mori-Akiyama et al. 2003) and *Six2*^-/-^ mice (Sweat et al. 2020). Sweat *et al*. further noted a direct up-stream regulatory relationship between the osteoprogenitor transcription factor, Six2, and Pax9, supporting a predicted role for Pax9 in progression of palate osteogenesis. The spatial transcriptomic alterations in *Pax9^-/-^* secondary palate cell clusters – specifically, the upregulation of Dkk2 surrounding the midline extension of palatal bone – provide evidence of Pax9’s functional role in regulating the differentiation and maturation of specific palatal mesenchymal progenitor cells. However, beyond the scope of the present work, more extensive functional studies on the connection between Pax9 and palatal mesenchymal osteoprogenitors in orchestrating their respective signaling pathways and cellular differentiation networks are needed.

The differential up-regulation of *Dkk1* and *Dkk2* within the *Pax9^-/-^* secondary palate mesenchyme is supportive of our previous foundational work in which these two molecules were targeted with small-molecule Wnt agonist therapeutics for the correction of embryonic cleft secondary palate (Jia et al. 2017a). Wnt signaling is known to directly influence *Runx2* transcription through its proximal promoter TCF binding site (Gaur et al., 2005). We hypothesize that the Wnt-antagonist *Dkk2* could indirectly reduce *Runx2 / Sp7* gene activation at the leading edge of the palatine bone, restricting expansion of this ossification center. It is unclear from the present study whether this is a driving or major component of *Pax9^-/-^* clefting. However, release of overall Wnt inhibition through *Wise* genetic deletion and Wnt-agonist treatment was sufficient to rescue the cleft palate, to varying extents (Jia et al., 2017; Li et al., 2017). Importantly, small molecule Wnt agonist IIIc3a’s inhibition of *Dkk1, Dkk2,* and *Dkk4* was 20% more effective to correct cleft palate *in utero* than WAY262611 inhibition of *Dkk1* alone (Jia et al., 2017).

In summary, this study addresses the perturbation of osteogenic patterning and differentiation in relation to Wnt signaling mediators in the *Pax9*^-/-^ genetic model of secondary cleft palate. We generated new multiomic and *in situ* spatial transcriptomic datasets of normal and abnormal palatogenesis to characterize the disrupted midline osteogenic extension of the palatine bone in the *Pax9*^-/-^ cleft secondary palate, alongside its perturbation in Wnt signaling dynamics, pointing to Pax9’s likely role in patterning and tuning Wnt-driven osteogenesis in the palate. We highlight for the first time, a spatial dysregulation of Wnt modulators and their overlapping pathway effectors in the *Pax9^-/-^* palate, specifically identifying *Dkk2* enriched in a boundary surrounding the osteogenic midline extension within palatal shelves of *Pax9^-/-^* mouse embryos. This contribution to our fundamental understanding of the molecular and cellular underpinnings of a cleft palate phenotype strengthens the framework required to assess the safety and efficacy of targeted therapeutic strategies to prevent or correct cleft palate disorders. Coupled with our prior proof-of-concept *in vivo* correction of cleft palate defects in *Pax9*^-/-^ mouse embryos, this additional knowledge of Wnt signaling perturbation allows for more precise preclinical testing of FDA-approved Wnt agonist therapies to correct such defects, both *in utero* and postnatally.

## Author Contributions

J.O. Piña contributed to conception, design, data acquisition and interpretation of data, performed all statistical analyses, drafted, and critically revised the manuscript; R. Raju contributed to conception, design, data acquisition, interpretation, and analysis, and critically revised the manuscript; D.M. Roth contributed to conception, design, and critically revised the manuscript; E.W. Winchester contributed to data acquisition, interpretation, and analysis, and critically revised the manuscript; C. Padilla contributed to data acquisition and critically revised the manuscript; J. Iben contributed to data acquisition and critically revised the manuscript; F.R. Faucz contributed to design, data acquisition and critically revised the manuscript; J.L. Cotney contributed to data acquisition, interpretation, and analysis, and critically revised the manuscript; R.N. D’Souza contributed to conception, design, interpretation of data, and critically revised the manuscript. All authors gave their final approval and agree to be accountable for all aspects of the work.

## Acknowledgements

We thank our lab manager, Parna Chattaraj, for her administrative support throughout manuscript preparation; lab member, Dr. Fahad Kidwai, for administrative oversight of animal colony logistics; Dr. Sergey Leiken (NIH/NICHD) for his technical and analytical support in establishing our independent next-generation sequencing workflow and Xenium *in situ* assays; Dr. Blake Warner (NIH/NIDCR) for his support in Xenium instrument access; Dr. Vardit Kram (NIH/NIDCR) for her guidance with µCT scanning and analysis; and Dr. Ricardo D. Coletta (University of Campinas, FOP, Brazil) for his review and feedback during manuscript preparation.

## Declaration of Conflicting Interests

The authors declared no potential conflicts of interest with respect to the research, authorship, and/or publication of this article.

## Funding

ZIA HD009015 “Wnt Signaling Pathway: Genes in Craniofacial Development: Palate and Tooth”, National Institutes of Health, National Institute of Dental and Craniofacial Research (RDS); National Institutes of Health, National Institute of Child Health and Human Development, Intramural Research Training Award (JOP/DMR); 1F30DE031149-01, 5R03DE028588-02, 2R35GM119465-06, 5R01DE028945-04 (JLC).

## Notes

### Competing Interest Statement

The authors have declared no competing interest.

### Summary of Updates

Novel highly-multiplex in situ mRNA localization profiling (Xenium In Situ, 10x Genomics, Inc.) for spatial resolution of palate marker genes using a custom gene panel derived from the multi-omic sequencing datasets generated herein.

