## Supplemental File for "Spatial Multiomics Reveal the Role of Wnt Modulator, Dkk2, in Palatogenesis"

| Appendix Table 1. Xenium In Situ custom panel gene list. | |
| --- | --- |
| name | ensembl_id |
| Hoatz | ENSMUSG00000032057 |
| Acan | ENSMUSG00000030607 |
| Acp5 | ENSMUSG00000001348 |
| Acta1 | ENSMUSG00000031972 |
| Acta2 | ENSMUSG00000035783 |
| Actr2 | ENSMUSG00000020152 |
| Actr3 | ENSMUSG00000026341 |
| Adamts17 | ENSMUSG00000058145 |
| Adipoq | ENSMUSG00000022878 |
| Ager | ENSMUSG00000015452 |
| Agtr1b | ENSMUSG00000054988 |
| Akt1 | ENSMUSG00000001729 |
| Alpl | ENSMUSG00000028766 |
| Alx1 | ENSMUSG00000036602 |
| Alx4 | ENSMUSG00000040310 |
| Amelx | ENSMUSG00000031354 |
| Angptl1 | ENSMUSG00000033544 |
| Anp32b | ENSMUSG00000028333 |
| Apoe | ENSMUSG00000002985 |
| Aqp5 | ENSMUSG00000044217 |
| Armc4 | ENSMUSG00000061802 |
| Atf4 | ENSMUSG00000042406 |
| Atf5 | ENSMUSG00000038539 |
| Atf6 | ENSMUSG00000026663 |
| Axin2 | ENSMUSG00000000142 |
| Bax | ENSMUSG00000003873 |
| Bcl11b | ENSMUSG00000048251 |
| Bglap | ENSMUSG00000074483 |
| Bgn | ENSMUSG00000031375 |
| Bicc1 | ENSMUSG00000014329 |
| Birc5 | ENSMUSG00000017716 |
| Bmp2 | ENSMUSG00000027358 |
| Bmp3 | ENSMUSG00000029335 |
| Bmp4 | ENSMUSG00000021835 |
| Bmp5 | ENSMUSG00000032179 |
| Bmp6 | ENSMUSG00000039004 |
| Bmp7 | ENSMUSG00000008999 |
| Bmpr1a | ENSMUSG00000021796 |
| Bmpr2 | ENSMUSG00000067336 |
| Brd2 | ENSMUSG00000024335 |
| Bst2 | ENSMUSG00000046718 |
| Calcrl | ENSMUSG00000059588 |
| Capsl | ENSMUSG00000039676 |
| Car2 | ENSMUSG00000027562 |
| Casq2 | ENSMUSG00000027861 |
| Ccl9 | ENSMUSG00000019122 |
| Ccn2 | ENSMUSG00000019997 |
| Ccn3 | ENSMUSG00000037362 |
| Ccn5 | ENSMUSG00000027656 |
| Cd200 | ENSMUSG00000022661 |
| Cd36 | ENSMUSG00000002944 |
| Cd63 | ENSMUSG00000025351 |
| Cd68 | ENSMUSG00000018774 |
| Cdh5 | ENSMUSG00000031871 |
| Cdk1 | ENSMUSG00000019942 |
| Cdkn1a | ENSMUSG00000023067 |
| Cenpf | ENSMUSG00000026605 |
| Cenpj | ENSMUSG00000064128 |
| Chad | ENSMUSG00000039084 |
| Chodl | ENSMUSG00000022860 |
| Cidea | ENSMUSG00000024526 |
| Ckm | ENSMUSG00000030399 |
| Cldn18 | ENSMUSG00000032473 |
| Cntn4 | ENSMUSG00000064293 |
| Col10a1 | ENSMUSG00000039462 |
| Col11a1 | ENSMUSG00000027966 |
| Col12a1 | ENSMUSG00000032332 |
| Col15a1 | ENSMUSG00000028339 |
| Col16a1 | ENSMUSG00000040690 |
| Col1a1 | ENSMUSG00000001506 |
| Col23a1 | ENSMUSG00000063564 |
| Col2a1 | ENSMUSG00000022483 |
| Col3a1 | ENSMUSG00000026043 |
| Col4a1 | ENSMUSG00000031502 |
| Crabp1 | ENSMUSG00000032291 |
| Creb3l1 | ENSMUSG00000027230 |
| Crocc2 | ENSMUSG00000084989 |
| Crp | ENSMUSG00000037942 |
| Crtap | ENSMUSG00000032431 |
| Csrp3 | ENSMUSG00000030470 |
| Ctsk | ENSMUSG00000028111 |
| Cxcl14 | ENSMUSG00000021508 |
| Dcn | ENSMUSG00000019929 |
| Ddit3 | ENSMUSG00000025408 |
| Deup1 | ENSMUSG00000039977 |
| Dio2 | ENSMUSG00000007682 |
| Dio3 | ENSMUSG00000075707 |
| Dkk1 | ENSMUSG00000024868 |
| Dkk2 | ENSMUSG00000028031 |
| Dkk4 | ENSMUSG00000031535 |
| Dlk1 | ENSMUSG00000040856 |
| Dlx3 | ENSMUSG00000001510 |
| Dlx5 | ENSMUSG00000029755 |
| Dlx6 | ENSMUSG00000029754 |
| Dmp1 | ENSMUSG00000029307 |
| Dspp | ENSMUSG00000053268 |
| Dthd1 | ENSMUSG00000090326 |
| Dvl2 | ENSMUSG00000020888 |
| Dynlrb2 | ENSMUSG00000034467 |
| Dysf | ENSMUSG00000033788 |
| Ebf2 | ENSMUSG00000022053 |
| Ecscr | ENSMUSG00000073599 |
| Egf | ENSMUSG00000028017 |
| Eif2ak1 | ENSMUSG00000029613 |
| Eif2ak2 | ENSMUSG00000024079 |
| Eif2ak3 | ENSMUSG00000031668 |
| Eif2ak4 | ENSMUSG00000005102 |
| Eif4ebp1 | ENSMUSG00000031490 |
| Eln | ENSMUSG00000029675 |
| Emcn | ENSMUSG00000054690 |
| Enam | ENSMUSG00000029286 |
| Eng | ENSMUSG00000026814 |
| Enpp2 | ENSMUSG00000022425 |
| Epha2 | ENSMUSG00000006445 |
| Ephb2 | ENSMUSG00000028664 |
| Erf | ENSMUSG00000040857 |
| Erg | ENSMUSG00000040732 |
| Esam | ENSMUSG00000001946 |
| Fat3 | ENSMUSG00000074505 |
| Fgf14 | ENSMUSG00000025551 |
| Fgf2 | ENSMUSG00000037225 |
| Fgf23 | ENSMUSG00000000182 |
| Fgf7 | ENSMUSG00000027208 |
| Fgfr1 | ENSMUSG00000031565 |
| Fgfr2 | ENSMUSG00000030849 |
| Fgfr3 | ENSMUSG00000054252 |
| Flt1 | ENSMUSG00000029648 |
| Fmod | ENSMUSG00000041559 |
| Fn1 | ENSMUSG00000026193 |
| Foxc1 | ENSMUSG00000050295 |
| Foxc2 | ENSMUSG00000046714 |
| Foxd2 | ENSMUSG00000055210 |
| Foxj1 | ENSMUSG00000034227 |
| Foxo3 | ENSMUSG00000048756 |
| Frem1 | ENSMUSG00000059049 |
| Frzb | ENSMUSG00000027004 |
| Fstl1 | ENSMUSG00000022816 |
| Fstl5 | ENSMUSG00000034098 |
| Gdf10 | ENSMUSG00000021943 |
| Gdf2 | ENSMUSG00000072625 |
| Gfra2 | ENSMUSG00000022103 |
| Gli1 | ENSMUSG00000025407 |
| Gli3 | ENSMUSG00000021318 |
| Gramd2 | ENSMUSG00000074259 |
| Grem1 | ENSMUSG00000074934 |
| Gsc | ENSMUSG00000021095 |
| Hmgn2 | ENSMUSG00000003038 |
| Hopx | ENSMUSG00000059325 |
| Hsp90ab1 | ENSMUSG00000023944 |
| Hspa5 | ENSMUSG00000026864 |
| Hspa9 | ENSMUSG00000024359 |
| Hspg2 | ENSMUSG00000028763 |
| Ibsp | ENSMUSG00000029306 |
| Icam1 | ENSMUSG00000037405 |
| Ifitm5 | ENSMUSG00000025489 |
| Igf1 | ENSMUSG00000020053 |
| Igf2 | ENSMUSG00000048583 |
| Igfbp4 | ENSMUSG00000017493 |
| Ihh | ENSMUSG00000006538 |
| Il6 | ENSMUSG00000025746 |
| Inha | ENSMUSG00000032968 |
| Inhba | ENSMUSG00000041324 |
| Itgav | ENSMUSG00000027087 |
| Itm2a | ENSMUSG00000031239 |
| Jag1 | ENSMUSG00000027276 |
| Jun | ENSMUSG00000052684 |
| Kif7 | ENSMUSG00000050382 |
| Krt1 | ENSMUSG00000046834 |
| Krt14 | ENSMUSG00000045545 |
| Krt5 | ENSMUSG00000061527 |
| Krt7 | ENSMUSG00000023039 |
| Lef1 | ENSMUSG00000027985 |
| Lfng | ENSMUSG00000029570 |
| Lgr5 | ENSMUSG00000020140 |
| Lif | ENSMUSG00000034394 |
| Lmnb1 | ENSMUSG00000024590 |
| Lrp5 | ENSMUSG00000024913 |
| Lrp6 | ENSMUSG00000030201 |
| Lrrc23 | ENSMUSG00000030125 |
| Ltbp2 | ENSMUSG00000002020 |
| Lum | ENSMUSG00000036446 |
| Map2k1 | ENSMUSG00000004936 |
| Mapk10 | ENSMUSG00000046709 |
| Mapk11 | ENSMUSG00000053137 |
| Mapk13 | ENSMUSG00000004864 |
| Mapk14 | ENSMUSG00000053436 |
| Mapk8 | ENSMUSG00000021936 |
| Matn4 | ENSMUSG00000016995 |
| Mdk | ENSMUSG00000027239 |
| Mfap2 | ENSMUSG00000060572 |
| Mgp | ENSMUSG00000030218 |
| Mki67 | ENSMUSG00000031004 |
| Mkx | ENSMUSG00000061013 |
| Mmp13 | ENSMUSG00000050578 |
| Mmp14 | ENSMUSG00000000957 |
| Mmp15 | ENSMUSG00000031790 |
| Mmp2 | ENSMUSG00000031740 |
| Mmp23 | ENSMUSG00000029061 |
| Mmp9 | ENSMUSG00000017737 |
| Mpz | ENSMUSG00000056569 |
| Mrc1 | ENSMUSG00000026712 |
| Msx1 | ENSMUSG00000048450 |
| Msx2 | ENSMUSG00000021469 |
| Mtor | ENSMUSG00000028991 |
| Myh8 | ENSMUSG00000055775 |
| Myl1 | ENSMUSG00000061816 |
| Mylk | ENSMUSG00000022836 |
| Myocd | ENSMUSG00000020542 |
| Nasp | ENSMUSG00000028693 |
| Ncam1 | ENSMUSG00000039542 |
| Ndrg2 | ENSMUSG00000004558 |
| Net1 | ENSMUSG00000021215 |
| Nfatc1 | ENSMUSG00000033016 |
| Nog | ENSMUSG00000048616 |
| Nos3 | ENSMUSG00000028978 |
| Notch1 | ENSMUSG00000026923 |
| Notch2 | ENSMUSG00000027878 |
| Nrg1 | ENSMUSG00000062991 |
| Ntrk1 | ENSMUSG00000028072 |
| Ogn | ENSMUSG00000021390 |
| Osr1 | ENSMUSG00000048387 |
| Palld | ENSMUSG00000058056 |
| Pax1 | ENSMUSG00000037034 |
| Pax9 | ENSMUSG00000001497 |
| Pcna | ENSMUSG00000027342 |
| Pdgfra | ENSMUSG00000029231 |
| Pdgfrb | ENSMUSG00000024620 |
| Pdpn | ENSMUSG00000028583 |
| Pecam1 | ENSMUSG00000020717 |
| Phactr1 | ENSMUSG00000054728 |
| Phex | ENSMUSG00000057457 |
| Piezo1 | ENSMUSG00000014444 |
| Piezo2 | ENSMUSG00000041482 |
| Podnl1 | ENSMUSG00000012889 |
| Postn | ENSMUSG00000027750 |
| Prickle1 | ENSMUSG00000036158 |
| Prrx1 | ENSMUSG00000026586 |
| Ptch1 | ENSMUSG00000021466 |
| Ptch2 | ENSMUSG00000028681 |
| Pth1r | ENSMUSG00000032492 |
| Pthlh | ENSMUSG00000048776 |
| Ptk2 | ENSMUSG00000022607 |
| Ptn | ENSMUSG00000029838 |
| Pxn | ENSMUSG00000029528 |
| Ran | ENSMUSG00000029430 |
| Ranbp1 | ENSMUSG00000005732 |
| Rfx1 | ENSMUSG00000031706 |
| Rgcc | ENSMUSG00000022018 |
| Rln1 | ENSMUSG00000039097 |
| Rock1 | ENSMUSG00000024290 |
| Rock2 | ENSMUSG00000020580 |
| Ror2 | ENSMUSG00000021464 |
| Rp1 | ENSMUSG00000025900 |
| Rspo1 | ENSMUSG00000028871 |
| Runx2 | ENSMUSG00000039153 |
| S100a6 | ENSMUSG00000001025 |
| Sapcd2 | ENSMUSG00000026955 |
| Scnn1g | ENSMUSG00000000216 |
| Selp | ENSMUSG00000026580 |
| Serpine1 | ENSMUSG00000037411 |
| Serpinf1 | ENSMUSG00000000753 |
| Serpinh1 | ENSMUSG00000070436 |
| Sfrp2 | ENSMUSG00000027996 |
| Sftpa1 | ENSMUSG00000021789 |
| Sftpc | ENSMUSG00000022097 |
| Sftpd | ENSMUSG00000021795 |
| Shh | ENSMUSG00000002633 |
| Six2 | ENSMUSG00000024134 |
| Slc2a4 | ENSMUSG00000018566 |
| Slit3 | ENSMUSG00000056427 |
| Smad1 | ENSMUSG00000031681 |
| Smad2 | ENSMUSG00000024563 |
| Smad3 | ENSMUSG00000032402 |
| Smad4 | ENSMUSG00000024515 |
| Smad5 | ENSMUSG00000021540 |
| Smad6 | ENSMUSG00000036867 |
| Smad7 | ENSMUSG00000025880 |
| Smad9 | ENSMUSG00000027796 |
| Smo | ENSMUSG00000001761 |
| Smurf1 | ENSMUSG00000038780 |
| Snai1 | ENSMUSG00000042821 |
| Sntn | ENSMUSG00000044772 |
| Sost | ENSMUSG00000001494 |
| Sostdc1 | ENSMUSG00000036169 |
| Sox11 | ENSMUSG00000063632 |
| Sox5 | ENSMUSG00000041540 |
| Sox6 | ENSMUSG00000051910 |
| Sox9 | ENSMUSG00000000567 |
| Sp7 | ENSMUSG00000060284 |
| Sparc | ENSMUSG00000018593 |
| Sparcl1 | ENSMUSG00000029309 |
| Spock1 | ENSMUSG00000056222 |
| Srf | ENSMUSG00000015605 |
| Stat3 | ENSMUSG00000004040 |
| Stat6 | ENSMUSG00000002147 |
| Stmn2 | ENSMUSG00000027500 |
| Sufu | ENSMUSG00000025231 |
| Tac1 | ENSMUSG00000061762 |
| Tbx1 | ENSMUSG00000009097 |
| Tbx15 | ENSMUSG00000027868 |
| Tcf12 | ENSMUSG00000032228 |
| Tek | ENSMUSG00000006386 |
| Tekt4 | ENSMUSG00000024175 |
| Tfap2a | ENSMUSG00000021359 |
| Tgfb1 | ENSMUSG00000002603 |
| Tgfb2 | ENSMUSG00000039239 |
| Tgfb3 | ENSMUSG00000021253 |
| Tgfbi | ENSMUSG00000035493 |
| Tgfbr3 | ENSMUSG00000029287 |
| Thbs1 | ENSMUSG00000040152 |
| Timp1 | ENSMUSG00000001131 |
| Timp2 | ENSMUSG00000017466 |
| Tln1 | ENSMUSG00000028465 |
| Tnf | ENSMUSG00000024401 |
| Tnfrsf11a | ENSMUSG00000026321 |
| Tnfrsf11b | ENSMUSG00000063727 |
| Tnfsf11 | ENSMUSG00000022015 |
| Tnik | ENSMUSG00000027692 |
| Tnn | ENSMUSG00000026725 |
| Top2a | ENSMUSG00000020914 |
| Trib3 | ENSMUSG00000032715 |
| Trnp1 | ENSMUSG00000056596 |
| Trpm3 | ENSMUSG00000052387 |
| Trps1 | ENSMUSG00000038679 |
| Tshr | ENSMUSG00000020963 |
| Ttn | ENSMUSG00000051747 |
| Tuba1b | ENSMUSG00000023004 |
| Tubb3 | ENSMUSG00000062380 |
| Twist1 | ENSMUSG00000035799 |
| Twist2 | ENSMUSG00000007805 |
| Tyrobp | ENSMUSG00000030579 |
| Unc5b | ENSMUSG00000020099 |
| Vcl | ENSMUSG00000021823 |
| Vdr | ENSMUSG00000022479 |
| Vim | ENSMUSG00000026728 |
| Vip | ENSMUSG00000019772 |
| Vwf | ENSMUSG00000001930 |
| Wif1 | ENSMUSG00000020218 |
| Wnt1 | ENSMUSG00000022997 |
| Wnt10a | ENSMUSG00000026167 |
| Wnt16 | ENSMUSG00000029671 |
| Wnt3a | ENSMUSG00000009900 |
| Wnt5a | ENSMUSG00000021994 |
| Wnt5b | ENSMUSG00000030170 |
| Wnt7a | ENSMUSG00000030093 |
| Wnt7b | ENSMUSG00000022382 |
| Wnt9a | ENSMUSG00000000126 |
| Xbp1 | ENSMUSG00000020484 |
| Zfp36 | ENSMUSG00000044786 |
| Zic1 | ENSMUSG00000032368 |

| Appendix Table 2. Quantitative transcript localization using Xenium in situ Analyzer. | | | | | | |
| --- | --- | --- | --- | --- | --- | --- |
|  | WT |  |  |  | PAX9^-/-^ |  |
|  | Transcripts |  |  |  | Transcripts |  |
| *Fgfr1* | 4165 |  |  |  | 11131 |  |
| *Gli1* | 498 |  |  |  | 1293 |  |
| *Gli3* | 613 |  |  |  | 1917 |  |
| *Bmp2* | 271 |  |  |  | 1003 |  |
| *Bmp4* | 191 |  |  |  | 395 |  |
| *Sfrp2* | 7561 |  |  |  | 18281 |  |
| *Dkk1* | 195 |  |  |  | 690 |  |
| *Ibsp* | 22736 |  |  |  | 40329 |  |
| *Phex* | 599 |  |  |  | 1006 |  |
| *Ptch1* | 1721 |  |  |  | 4650 |  |
| *Matn4* | 114 |  |  |  | 791 |  |
| *Mmp9* | 615 |  |  |  | 285 |  |
| *Mmp13* | 695 |  |  |  | 110 |  |
| *Mmp14* | 4082 |  |  |  | 9009 |  |
| *Mki67* | 2648 |  |  |  | 9043 |  |
| *Dkk2* | 375 |  |  |  | 975 |  |
|  | Area [µm^2^] | 1275539.3 |  |  | Area [µm^2^] | 1273533.9 |
|  | Cells | 9320 |  |  | Cells | 10647 |

| Appendix Table 3. Xenium In Situ quality output for all 6 samples analyzed. | | |
| --- | --- | --- |
| region_name | panel_name | region_area |
| PAX9_KO_1 | 7GUZV9_mOther_350g_gene_panel | 22338455.36 |
| PAX9_KO_2 | 7GUZV9_mOther_350g_gene_panel | 20591083.78 |
| PAX9_KO_3 | 7GUZV9_mOther_350g_gene_panel | 13171462.87 |
| WT1 | 7GUZV9_mOther_350g_gene_panel | 25823386.6 |
| WT2 | 7GUZV9_mOther_350g_gene_panel | 20589879.64 |
| WT3 | 7GUZV9_mOther_350g_gene_panel | 17976017.34 |
| region_name | total_cell_area | fraction_transcripts_decoded_q20 |
| PAX9_KO_1 | 15856693.13 | 0.879755214 |
| PAX9_KO_2 | 14587457.78 | 0.879952086 |
| PAX9_KO_3 | 9495038.628 | 0.87233069 |
| WT1 | 17924500.31 | 0.872153274 |
| WT2 | 15141570.23 | 0.882587815 |
| WT3 | 14421495.82 | 0.885747601 |
| region_name | fraction_empty_cells | num_cells_detected |
| PAX9_KO_1 | 9.93E-05 | 171167 |
| PAX9_KO_2 | 0.000111553 | 161358 |
| PAX9_KO_3 | 0.000214794 | 88457 |
| WT1 | 0.000329298 | 154875 |
| WT2 | 4.96E-05 | 141190 |
| WT3 | 0 | 145244 |
| region_name | decoded_transcripts_per_100um2 | adjusted_negative_control_probe_rate |
| PAX9_KO_1 | 93.08025248 | 0.001402373 |
| PAX9_KO_2 | 91.85441495 | 0.001838757 |
| PAX9_KO_3 | 92.5039312 | 0.003745976 |
| WT1 | 54.78866821 | 0.001533184 |
| WT2 | 74.18860677 | 0.00116255 |
| WT3 | 99.23720242 | 0.001223978 |
| region_name | adjusted_negative_control_codeword_rate | fraction_transcripts_assigned |
| PAX9_KO_1 | 0.000285145 | 0.997097792 |
| PAX9_KO_2 | 0.000252059 | 0.997473361 |
| PAX9_KO_3 | 0.000355942 | 0.997543288 |
| WT1 | 0.000286148 | 0.991514465 |
| WT2 | 0.000239596 | 0.994286106 |
| WT3 | 0.000294995 | 0.996995002 |
| region_name | median_genes_per_cell | median_transcripts_per_cell |
| PAX9_KO_1 | 35 | 71 |
| PAX9_KO_2 | 35 | 69 |
| PAX9_KO_3 | 42 | 88 |
| WT1 | 27 | 52 |
| WT2 | 33 | 68 |
| WT3 | 38 | 86 |

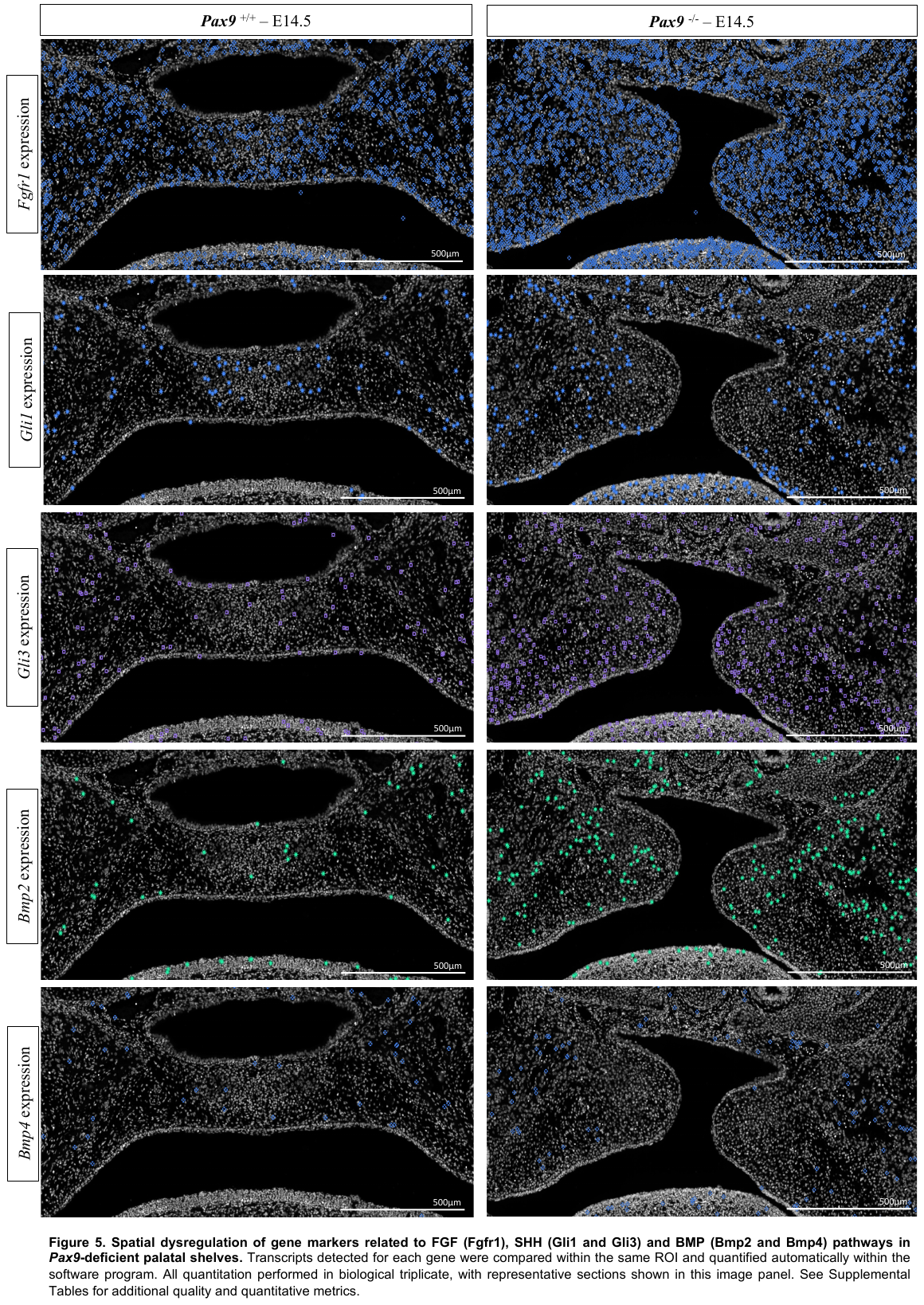
